# Molecular dynamics simulations of human cohesin subunits identify DNA binding sites and their potential roles in DNA loop extrusion

**DOI:** 10.1101/2024.09.17.613402

**Authors:** Chenyang Gu, Shoji Takada, Giovani B. Brandani, Tsuyoshi Terakawa

## Abstract

The SMC complex cohesin mediates interphase chromatin structural formation in eukaryotic cells through DNA loop extrusion. Here, we sought to investigate its mechanism using molecular dynamics simulations. To achieve this, we first constructed the amino-acid-residue-resolution structural models of the cohesin subunits, SMC1, SMC3, STAG1, and NIPBL. By simulating these subunits with double-stranded DNA molecules, we predicted DNA binding patches on each subunit and quantified the affinities of these patches to DNA using their dissociation rate constants as a proxy. Then, we constructed the structural model of the whole cohesin complex and mapped the predicted high-affinity DNA binding patches on the structure. From the spatial relations of the predicted patches, we identified that multiple patches on the SMC1, SMC3, STAG1, and NIPBL subunits form a DNA clamping patch group. The simulations of the whole complex with double-stranded DNA molecules suggest that this patch group facilitates DNA bending and helps capture a DNA segment in the cohesin ring formed by the SMC1 and SMC3 subunits. In previous studies, these have been identified as critical steps in DNA loop extrusion. Therefore, this study provides experimentally testable predictions of DNA binding sites implicated in previously proposed DNA loop extrusion mechanisms and highlights the essential roles of the accessory subunits STAG1 and NIPBL in the mechanism.

## Introduction

Cohesin, a structural maintenance of chromosomes (SMC) complex, is a molecular motor that mediates chromosome structural regulation in eukaryotic cells. In addition to its role in sister chromatid cohesion, cohesin is involved in the formation of topologically associating domain (TAD) in interphase chromatin (1): self-interacting genomic regions insulated from the outside (2). TAD formation plays a critical role in gene expression regulation by facilitating or preventing the interactions between promoters and enhancers (3,4). As a mechanism for the TAD formation by cohesin, the DNA loop extrusion mechanism (5) was previously proposed. During loop extrusion, cohesin captures a small DNA segment inside its ring-like protein structure and expands it by extrusion until it encounters insulator elements such as CTCF. The mechanism was proposed based on the observation that cohesin and pairs of convergent CTCF binding sites are often co-localized at the boundaries of TADs (6). Single-molecule imaging studies confirmed that cohesin, as a molecular motor, can extrude a DNA loop using adenosine triphosphate (ATP) hydrolysis energy (7,8).

The human cohesin complex forms a hetero-pentamer (Fig 1). The SMC1 and SMC3 subunits dimerize at their hinge domain and can also engage at their ATPase head domain, depending on the bound nucleotides. The N- and C-terminal regions of the intrinsically disordered RAD21 subunit attach to the SMC3 and SMC1 head domains, respectively. The two accessory subunits, STAG1 and NIPBL, bind to RAD21 and are essential for motor activity, though their exact roles still need to be clarified (7,8). Cohesin changes its conformation or DNA-binding state throughout the ATP hydrolysis cycle (9–11). SMC1/3 heads engage upon ATP binding and disengage after ATP hydrolysis, while NIPBL associates with RAD21 upon the head engagement and dissociates after ATP hydrolysis. Upon ATP binding to SMC1/3, about 50 nm-long anti-parallel coiled-coil arms that connect their heads and hinges are curved to take an open “O” conformation (Fig 1A). The entire cohesin complex transitions to an “8”-like conformation (Fig 1B) upon the head engagement, with the whole complex forming two distinct compartments where DNA could potentially be entrapped: one formed by the coiled coils from the hinge up to the engaged heads, and another formed by RAD21. Upon head disengagement, these two compartments merge into one larger compartment. These conformational changes have been suggested to drive DNA loop extrusion (12,13).

**Fig 1.**
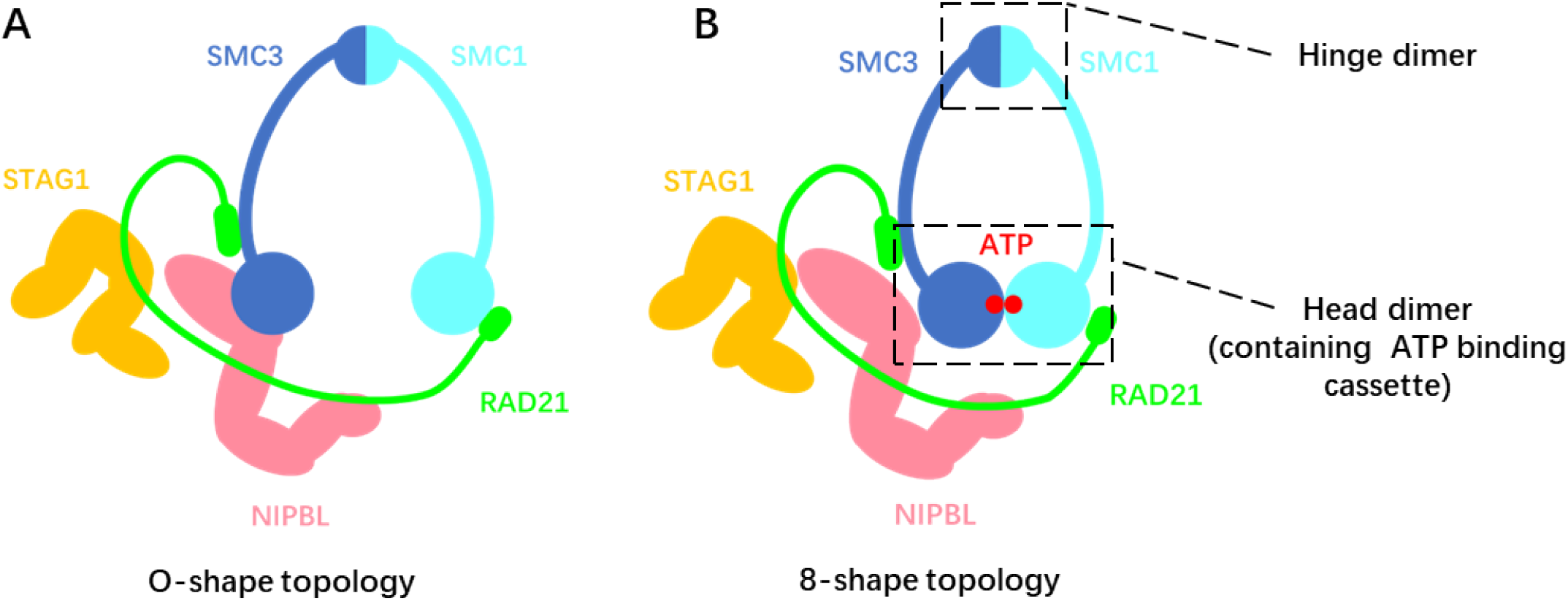
An illustration of the human cohesin complex in (A) O-shape and (B) 8-shape topology.

Previous studies have adopted numerous experiments, including high-speed atomic force microscopy (HS-AFM) (9,10), cryogenic electron microscopy (cryo-EM) (14,15), and single-molecule fluorescence imaging (7,8), to investigate the interaction between SMC complexes and DNA. Despite their success, HS-AFM and fluorescence imaging need more resolution to identify the precise DNA-binding sites. Cryo-EM requires averaging over many structural snapshots to obtain high-resolution structures, potentially missing temporary interactions between DNA and other parts of the complex. For example, mutating the basic amino acids on the surface of SMC1/3 hinge dimer facing the inside of the ring compartment formed by coiled coils or on the C-terminus of STAG1’s HEAT repeats domain abrogate loop extrusion (9), but DNA binding at these sites was not observed by cryo-EM. Therefore, it is reasonable to assume that some critical DNA binding sites involved in loop extrusion have not been resolved in the previous works. So far, multiple models for the loop extrusion mechanism have been proposed, such as the DNA segment capture model (13), the scrunching model (5), the swing and clamp model (9), and the hold and feed model (16). They all assume that a DNA-binding site captures a DNA segment and hands it over to another throughout the ATP hydrolysis cycle. Therefore, identifying new DNA binding sites is essential for validating and refining these models or proposing new ones. To date, Higashi et al. (17) and Nomidis et al. (18) have performed 3D simulations using coarse-grained models that represent cohesin or SMC complexes in general, respectively, using a small number of rigid segments connected by flexible joints. Both studies have obtained valuable insights into the DNA loop extrusion mechanism. However, the models omitted or highly coarse-grained the interactions between protein and DNA. Prediction of DNA binding sites should help refine such coarser models.

In this study, we performed molecular dynamics (MD) simulations of human cohesin subunits with DNA to comprehensively predict and rank the strength of the DNA binding sites on these proteins. We also modeled the whole cohesin complex and mapped the predicted binding site to infer the potential pathways of DNA handover in the context of DNA loop extrusion.

## Result

### Residue-resolution modeling of human cohesin subunits

In this work, we predicted DNA binding sites in cohesin subunits and quantified their strength by measuring the rates of DNA unbinding from the simulations. For this computationally demanding task, we used the AICG2+ protein model, where one particle represents one amino acid, and the 3SPN.2C DNA model (19,20), where three particles represent one nucleotide. Debye-Hückel electrostatic interactions and excluded volume interactions were imposed between protein and DNA. This model allows a speedup of several orders of magnitude compared to all-atom models while retaining key molecular details and predictive capacity, having been already applied to a variety of systems, including *S. cerevisiae* condensin (21), human PRC2 complex (22), and DNA binding proteins HoxD9, Sap1, and Skn1 (23), to predict their DNA binding sites successfully.

We built residue-resolution coarse-grained structural models of essential subunits of human cohesin. The initiation and progression of DNA loop extrusion require the SMC1, SMC3, RAD21, and STAG1 subunits and the C-terminal region of the NIPBL subunit in addition to ATP and DNA (7,8). We chose these five subunits as our target. The cryo-EM structure of the human cohesin-DNA complex (PDB ID: 6WG3) (14) shows the gripping state in which these subunits load on DNA with the heads of SMC1 and SMC3 bound to ATP. We used the partial structures from this complex to build the residue-resolution structural model of each subunit.

The cryo-EM structure has missing residues: the parts of the coiled-coil arms in SMC1 and SMC3 and the intrinsically disordered regions (IDR) flanking the HEAT repeats of STAG1 and NIPBL. Therefore, we built residue-resolution structural models of the following eight constructs (Fig 2A to H): (A) the SMC1 head and (B) the SMC3 head with the cryo-EM solved part of the coiled-coil arms, (C) the dimer of A and B, (D) the SMC1 hinge, (E) the SMC3 hinge, (F) the dimer of D and E, (G) the HEAT repeats of STAG1, and (H) NIPBL. These include monomeric and heterodimeric domains to investigate the cooperatively created binding site. We omitted the IDR of STAG1 and NIPBL from the models because of their low sequence conservation, suggesting a minor contribution to loop extrusion (Fig S1). Consistent with this, the N-terminal IDR of NIPBL has been proven unessential for *in vitro* DNA loop extrusion (8), while the C-terminal IDR of STAG1 and NIPBL plays a role in loading to DNA (10).

**Fig 2.**
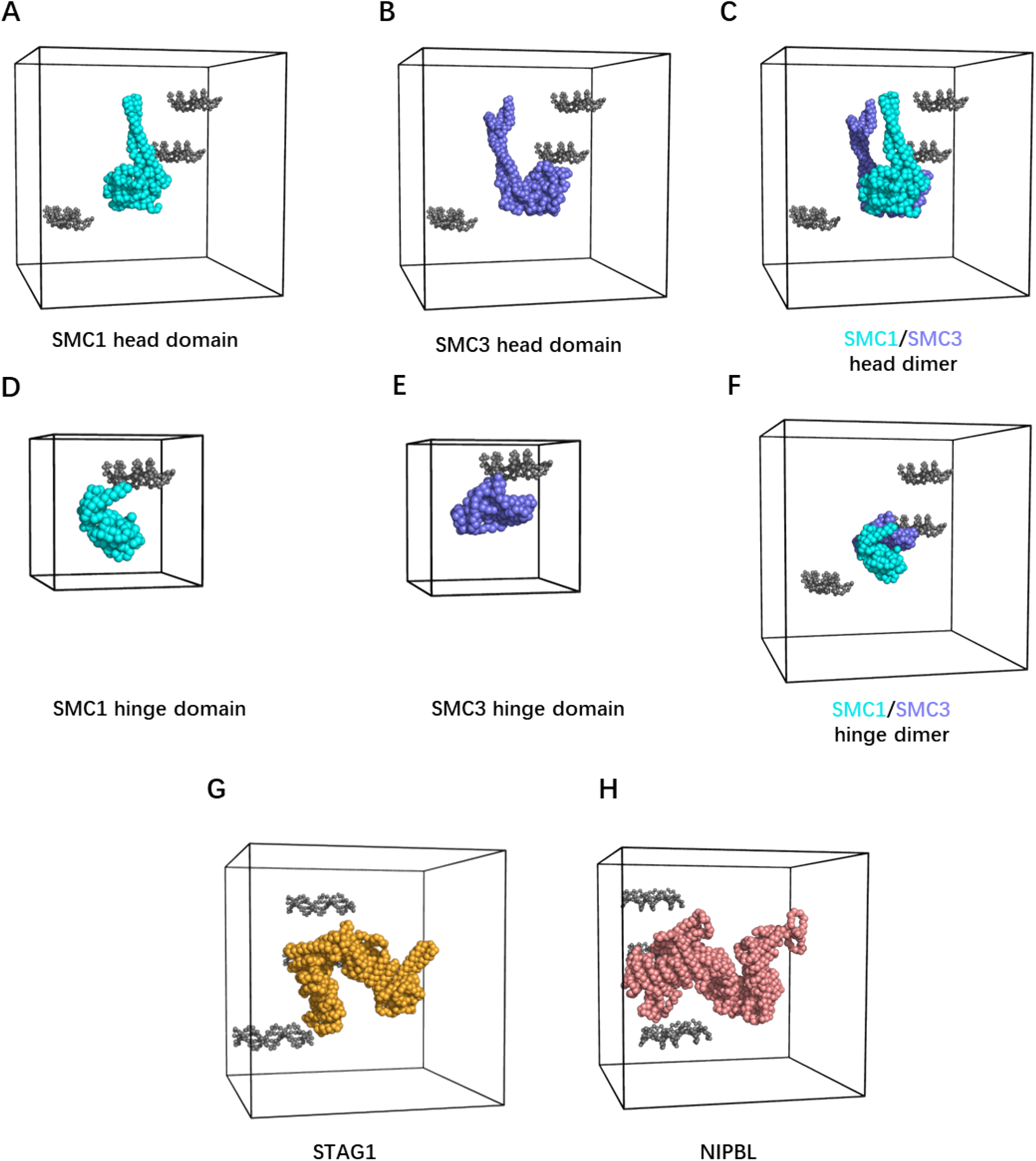
Residue-resolution modelling of human cohesin subunits and DNA binding simulation setup. (A to H) Simulation setup of DNA binding assays for cohesin subunits. Gray-colored beads indicate the DNA model, and the black cube indicates the periodic simulation boundary. Cyan, blue, orange, and pink colored beads indicate SMC1 head domain with an emanating partial coiled-coil arm in (A), SMC3 head domain with an emanating partial coiled-coil arm in (B), SMC1/3 head dimer with emanating partial coiled-coil arms in (C), SMC1 hinge domain in (D), SMC3 hinge domain in (E), SMC1/3 hinge dimer in (F), the HEAT repeats domain of STAG1 in (G), and the HEAT repeats domain of NIPBL in (H).

We then prepared residue-resolution models of short DNA segments. In the cryo-EM structure (14), cohesin forms a complex with 72 base pairs (bp) of double-stranded DNA. However, the experiment only captures 17 bp of DNA bound to cohesin and does not identify the sequence of the captured DNA. To study the non-specific DNA binding of the cohesin subunits, we used a randomly generated 20 bp DNA sequence with the same GC content as the DNA used in the cryo-EM experiment (14) (Methods). When multiple DNA segments were introduced, we used copies of the same DNA sequence.

The residue-resolution structural models of each protein subunit and DNA segments were constructed using CafeMol (24) (Fig 2). In the AICG2+ protein model and forcefield (25), one particle at the α-carbon position represents one amino acid residue. The force field stabilizes the native protein conformation, with the equilibrium bond lengths, bond angles, dihedral angles, and non-bonded native contact lengths extracted from reference all-atom structures. In the 3SPN.2C DNA model (20), three particles at the centers of mass of the base, sugar, and phosphate units represent one nucleotide. Proteins and DNA interact via Debye-Hückel electrostatics, which may cause attraction/repulsion, and excluded volume, which is purely repulsive. Partial charges on the protein particles were arranged using the RESPAC algorithm (26) to reproduce the electrostatic potential around the reference all-atom structures.

We performed residue-resolution coarse-grained MD simulations to identify DNA-binding sites on cohesin subunits and quantify the affinities (Fig 2A to H). Protein structural models SMC1 head, SMC3 head, SMC1/3 head dimer, SMC1/3 hinge dimer, NIPBL, and STAG1 are each placed inside a small cubic box with periodic boundaries, together with three copies of 20 bp DNA fragments randomly placed around the protein (see Methods section for more details). Including multiple DNAs increases the efficiency of the binding site search thanks to the high concentration and the fact that two DNA segments can continue exploring even if one is trapped at a particular site. SMC1 hinge and SMC3 hinge are each placed inside a periodic box with only one copy of the 20 bp DNA since the small size of these subunits naturally facilitates the identification of the binding sites.

### Molecular dynamics simulations predicted DNA binding patches in cohesin

We performed residue-resolution MD simulations on systems A to H as described above (Movie 1). In all systems (using system A, SMC1 head domain with short emanating coiled coils, as an example in Fig 3), the simulation trajectories showed DNA segments repeatedly associating with and dissociating from the protein surface residues (Fig 3A). We considered a DNA segment in contact with an amino acid when the smallest distance from any particle of the DNA segment to the amino acid particle is within a given threshold (see Methods for details). Based on this definition, we identified the amino acids in contact with each DNA segment as a function of time (Fig 3B). Surface amino acids that are spatially close to each other tend to encounter the same DNA segment at a given time, and DNA-contacting probabilities mapped on the structure show that amino acid particles with high contact probability cluster at several spatially localized regions (Fig S2). Hereafter, we refer to these spatial regions as binding patches. The clear pattern of simultaneous binding with DNA within each patch allows us to quantitatively identify patches with a clustering analysis using the Jaccard index describing the similarity in DNA contact patterns between two amino acid particles (Methods, Fig S3). The identified binding patches are mapped on the structure (Fig 3C, Fig S4).

**Fig 3.**
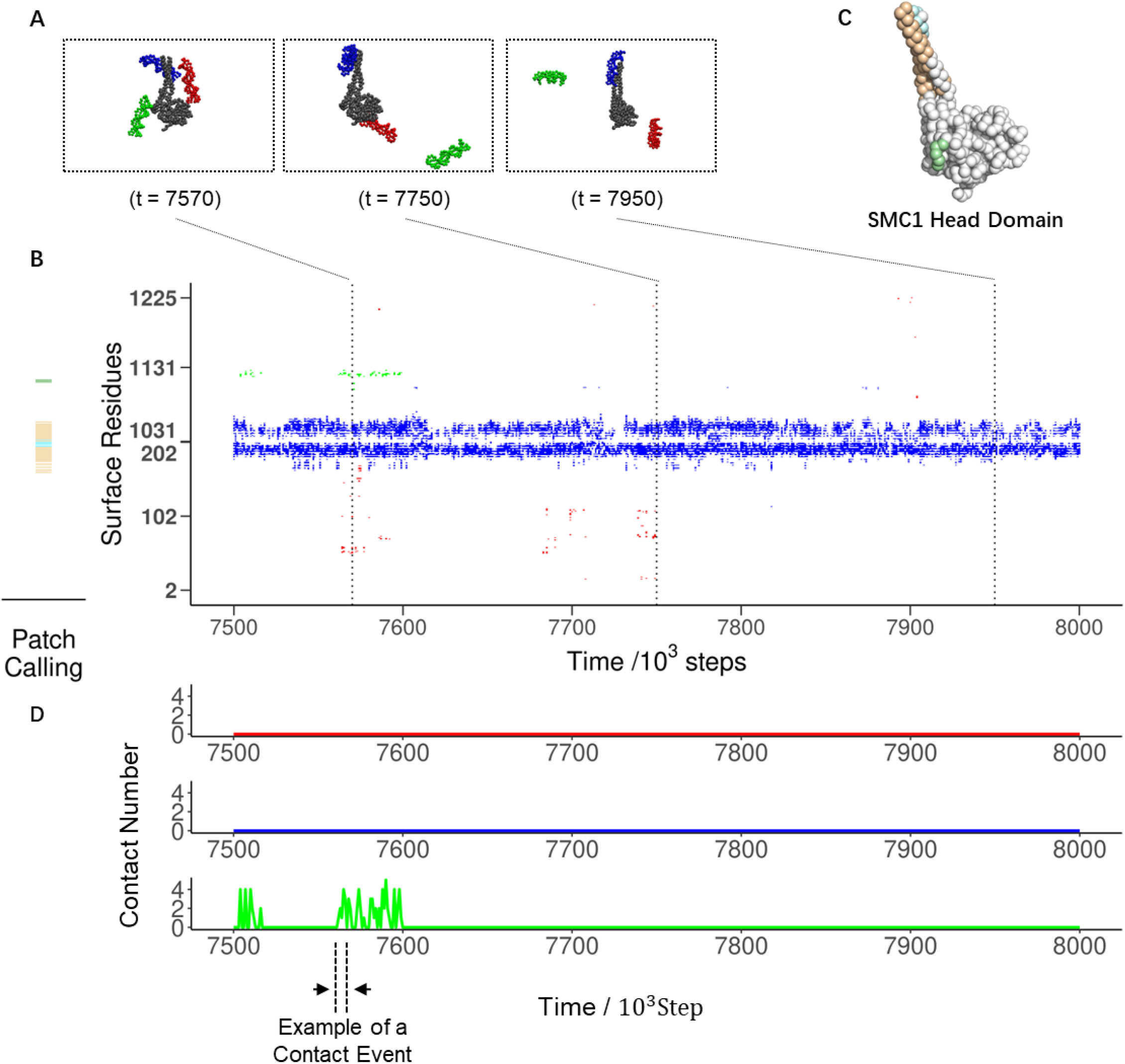
Analysis of DNA binding simulation assay trajectories. (A) Representative snapshots of the simulation of the SMC1 head domain with DNA (System A in Fig 2) taken from the trajectory shown in (B). The numbers indicate simulation time step *t* in the unit of 1000 steps. One (right), two (center), or three (left) DNA segments are in contact with SMC1 surface residues in each snapshot. (B left) DNA binding patches identified on the SMC1 head domain using all 20 trajectories. (B right) Time series of IDs of amino acid particles in contact with DNA. The colors indicate which DNA (red, blue, or green) in (A) is in contact with the protein. (C) Binding patches presented in (B left) mapped on the structure of the SMC1 head domain. (D) Time series of the number of DNA-contacting residues in the light green patch in (C). An example of a single contact event is shown with dotted lines and arrows.

To evaluate the DNA-binding affinity of each patch, we calculated their dissociation rate constants from the simulation trajectories. We reasonably assumed that DNA binding is diffusion-limited, implying a similar association rate constant for all patches, which allows us to use the dissociation rate constant as a proxy for relative affinity. To estimate the dissociation rates, we measured the time intervals in which DNA stays in contact with each patch [Fig 3D, using patch A2 as an example. A2 is colored light green in Fig 3B left panel and Fig 3C. Throughout this paper, we systematically name patches by combining the name of the system (here A) and the affinity rank (here second).] and used these to compute their survival probabilities (Fig S5, Methods), representing the fraction of DNA molecules staying on the patch after a particular duration. Fitting exponential functions to the curve provided us with the dissociation rate constants. The sum of two exponential functions fitted the curves markedly better than one exponential function for all the patches (Fig S5), suggesting the two interaction modes: the transient and stable modes. Here, we used the dissociation rate constant of the stable mode (the lower one) as a proxy for relative affinity to ignore the random encounter.

The analysis identified multiple DNA binding patches on each subunit and its dimeric form (Table 1). We focused on the patches that associate more strongly with DNA than the patch on the top surface of the heads of SMC1/3 heterodimer (Fig 4A), as those most likely to play a significant role in loop extrusion, as DNA is clamped on this surface in the cryo-EM structure of the gripping state (14). Therefore, we predicted the DNA binding patches with a dissociation rate constant lower than 2.0 × 10^―5^timesteps^-1^ (Fig 4B to E) as the potential DNA binding patches. Table 1 lists these binding patches, their dissociation rate constant, key positively charged residues, and eventual remarks on the patches.

**Table 1.** DNA Binding patches with a binding affinity strong enough to participate in DNA handover during translocation or loop extrusion.

| Subunit/<br>domain | patch<br>name | $k_{off}$ | Positive charged residues | Remarks |
| --- | --- | --- | --- | --- |
| **SMC1**<br>**head** | A1 | $1.36 \times 10^{-7}$ | 170, 171, 187, 188, 189, 190, 196, 197,<br>200, 1049, 1050, 1051, 1053, 1054, 1056,<br>1064 | |
| **SMC3**<br>**head** | B1 | $1.26 \times 10^{-5}$ | 33, 55, 57, 143, | |
| | B2 | $1.63 \times 10^{-5}$ | 72, 105, 106, 113, 185, 997, 1003 | |
| **SMC1/3**<br>**head**<br>**dimer** | C1 | $1.26 \times 10^{-7}$ | SMC1: 170, 171, 172, 173, 187, 188, 189,<br>190, 196, 197, 200, 1035, 1049, 1050,<br>1051, 1053, 1054, 1056 | ① |
| | C2 | $1.56 \times 10^{-5}$ | SMC1: 59 | ② |
| | C3 | $1.62 \times 10^{-5}$ | SMC3: 55, 57, 72, 105, 106, 113, 114 | ③ |
| | C4 | $1.64 \times 10^{-5}$ | SMC3: 185, 188, 997, 999, 1002, 1003, | |
| **SMC1/3**<br>**hinge**<br>**dimer** | F1 | $2.18 \times 10^{-6}$ | SMC1: 491, 496, 499, 500, 554, 561, 564;<br>SMC1: 493, 498, 503, 514, 518, 520, 521,<br>522, 527, 533, 557, 561, 571, 578, 592,<br>596, 612, 614, 618, 621, 624, 625, 629,<br>634, 644, 660, 661, 672, 673, 675, 680,<br>683 | |
| **STAG1** | G1 | $5.59 \times 10^{-6}$ | 172, 173, 175, 352, 549, 550 | |
| | G2 | $7.03 \times 10^{-6}$ | 718, 721, 759, 766, 767, 770, | |
| | G3 | $9.60 \times 10^{-6}$ | 385, 910, 912, 957, 958, 1013, 1016 | |
| **NIPBL** | H1 | $3.37 \times 10^{-7}$ | 1235, 1236, 1247, 1294, 1297, 1369, 1371,<br>1372, 1374, 1378, 1379, 1381, 1389 | ④ |
| | H2 | $9.32 \times 10^{-7}$ | 1552, 1599, 1789, 1791, 1794, 1828, 1865,<br>1866, 1867, 1870, 1873, 1904, 1912 | ⑤ |
| | H3 | $5.43 \times 10^{-6}$ | 2104, 2112, 2119, 2162, | |
| | H4 | $7.71 \times 10^{-6}$ | 1779, 2563, 2574, 2578, 2583 | |
| | H5 | $1.02 \times 10^{-5}$ | 1965, 1968, 1969 | |

**Fig 4.**
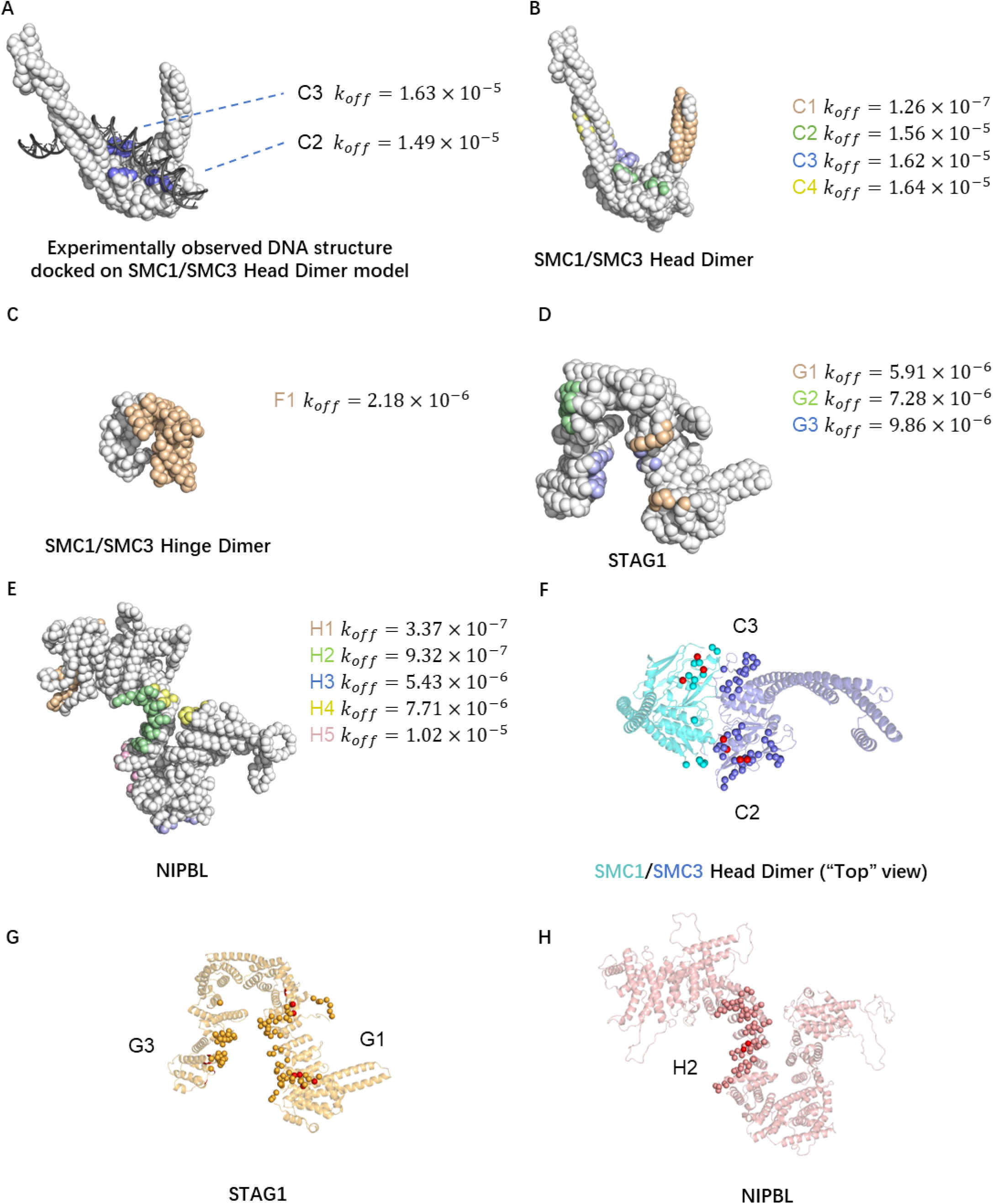
Prediction and validation of strong DNA binding patches. (A) The positions of DNA binding patches on the top surface of the SMC1/3 head dimer. The cryo-EM DNA structure (PDB ID: 6WG3) is superimposed on the coarse-grained model of the SMC1/3 head dimer. (B to E) DNA binding patches with a binding affinity higher or comparable with patches in (A) on each subunit or their dimers. (F to H) Positions of the predicted strong patches (beads colored other than red) and experimentally identified critical amino acid residues (red beads) in the SMC1/3 head dimer, STAG1, and NIPBL.

In the SMC1 head domain (A) and head dimer systems (C), we observed strong patches A1 and C1 (Table 1), corresponding to the same group of amino acids on the coiled-coil part connected to the SMC1 head domain. These patches always bind to at least one DNA segment throughout the simulation trajectories. This means the dissociation of one DNA segment from these patches results from another DNA segment competing for the same binding site (Fig S6). Although association duration for these patches still follows exponential distributions, the dissociation rate constants calculated under this condition have different physical meanings than other patches and should not be compared directly. However, the value we obtained in this way gives us a lower bound on the ideal dissociation rate constants. Since the values for these patches are the smallest among all the identified patches, we know that these patches have strong interaction with DNA and should be considered essential candidates as patches involved in DNA loop extrusion.

In each system, multiple DNA binding patches are identified (Fig S4, Table 1). In the dimer systems SMC1/3 head dimer (C) and SMC1/3 hinge dimer (F), dimerization caused some surface amino acids to be hidden, altering the DNA binding patches in the corresponding monomers. The dimerization partly hides patch B1 (side surface of SMC3 head domain), while the exposed part forms a new patch C3 (top surface of SMC1/3 head dimer) with some amino acids on the SMC1 head domain (Fig S4A to C). Dimerization of the SMC1 and SMC3 hinge domains unifies the patches D2, E1, E2, E3, and E4. These patches collectively form a patch F1, which contains all the surface residues of the SMC3 hinge domain and some amino acids from the SMC1 hinge domain (Fig S4D to F). SMC1 and SMC3 head domains engage and disengage during the ATP hydrolysis cycle powering DNA loop extrusion, making dynamic changes in DNA binding patches potentially relevant for the loop extrusion mechanism. Hinge opening is also hypothesized to play a role in the loading process of cohesin (27). Therefore, binding patches on SMC1 head domain (A), SMC3 head domain (B), SMC1 hinge domain (D), SMC3 hinge domain (E), SMC1/3 head dimer (C), and SMC1/3 hinge dimer (F) should all be investigated when studying the loop extrusion mechanism.

### Electrostatics and previous mutagenesis validated the predicted DNA-binding patches

We now turn to the role of electrostatic interactions in the identified DNA-binding patches on the proteins. In this study, the RESPAC algorithm determined charge arrangements on surface amino acid particles to reproduce the electrostatic potential around an all-atom protein structure calculated by the Adaptive Poisson-Boltzmann Solver (28). The positive electrostatic potential surface area overlapped the predicted DNA-binding patches in the simulation runs. Notably, when we replaced the RESPAC charges with simpler ±1 unit charges on basic and acidic amino acid particles, the DNA binding sites appeared more dispersed, thus weakening some key electrostatic features (e.g. the strong and concentrated positive charges at the ‘neck’ of NIPBL is recreated by RESPAC calibration, Fig S7). Therefore, the charge arrangement calibration by the RESPAC algorithm assisted in predicting the DNA binding sites created by the cooperative contribution of charges.

The predicted DNA binding patches also overlap with the amino acid residues already identified to play an essential role in DNA loop extrusion in the previous mutagenesis studies (9) (Fig 4F to H). Bauer *et al*. showed that mutations or deletions of some surface residues of SMC1/3, STAG1, or NIPBL abrogate or slow down loop extrusion (Table 2). These residues shown in Fig 4F to H were in the predicted DNA-binding sites, validating our simulation result.

**Table 2.** Predicted DNA binding patches that spatially or sequentially overlap with the experimentally identified critical amino acids.

| Simulation predicted patch | Experiment | Effect on loop extrusion |
| --- | --- | --- |
| Head dimer patch #2 | SMC1 K52E, R57E, K59E, K62E | Abrogate |
|  | ΔSMC1 58-62 | Reduce |
| Head dimer patch #3 | SMC3 R57E, R61E, K105E, K106E | Abrogate |
|  | SMC3 R57E, R61E | Reduce |
| STAG1 patch #1 | STAG1 K92E, K95E, K172E, K173E | Abrogate |
|  | STAG1 K555E, K558E, K564E | Abrogate |
| STAG1 patch #3 | STAG1 K969E, K971E, K1013E, K1016E | Abrogate |
| NIPBL patch #2 | NIPBL R1867A, R1870A | Abrogate |

Our simulations also predicted new DNA-binding patches, including residues that have yet to be considered in mutagenesis studies. For example, we identified a potential DNA binding site at the coiled-coil arm emanating from the SMC1 head. The crystal structure of Rad50 in complex with DNA supported this prediction (29). Rad50 is a DNA repair protein paralogous to cohesin and shares a similar amino acid sequence and molecular architecture. In the structure, DNA binds to the coiled-coil arm region emanating from the head, whose sequence is conserved to the corresponding predicted DNA binding site on cohesin and located at a similar location (Fig S8). Therefore, our MD simulations predicted experimentally testable potential DNA binding patches.

### Cooperative contributions of DNA binding sites may drive loop extrusion

We conducted structural modeling of the whole cohesin complex, including SMC1, SMC3, STAG1, NIPBL, and RAD21, to get insight into the relative positions of the DNA binding sites predicted above and to infer the possible pathways of DNA handover in DNA loop extrusion (Fig 5A).

**Fig 5.**
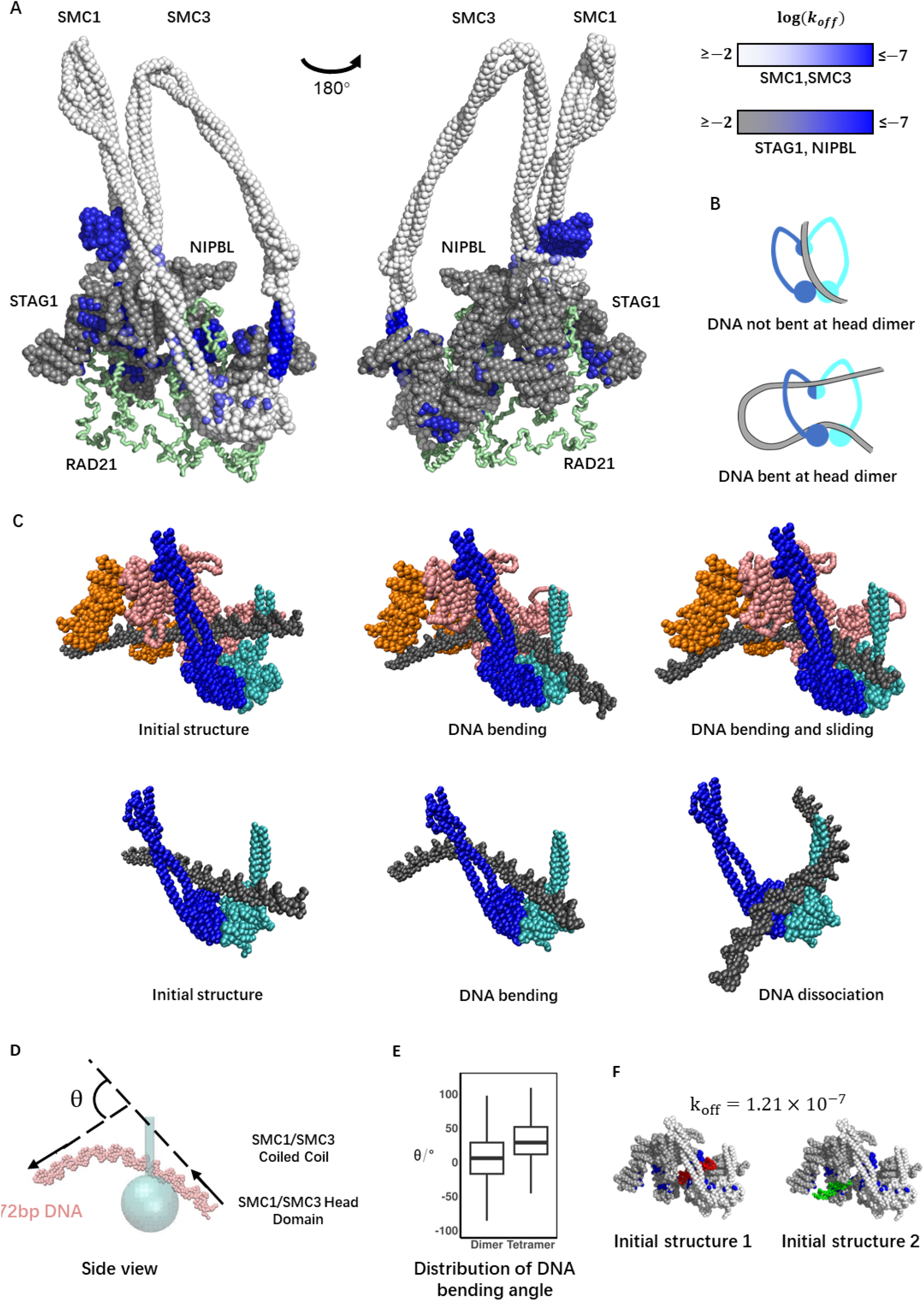
Spatial relations of DNA binding patches in the whole cohesin complex and the cooperative effect of the patch group. (A) The positions of all significant DNA binding patches identified in Fig 4 mapped on the structural model of the whole cohesin complex. DNA binding affinity is indicated by blue intensity. The amino acid particles in SMC1 and SMC3 not identified as significant patches are colored in white, while those in STAG1 and NIPBL are colored in gray. Rad21 is visualized as green tube. (B) Schematics of the DNA segment capture model. (C) The setup and representative snapshots of the simulations studying the effect of the DNA-gripping patch group. (C top) The simulation of the tetrameric complex containing the DNA-gripping patch group (SMC1/3 heads, STAG1, and NIPBL) with 72 bp dsDNA. (C bottom) The simulation of the SMC1/3 heads and the coiled coils emanating from them containing the partial DNA-gripping patch group (SMC1/3 heads). (D) An example snapshot of DNA projected on the plane cutting the cohesin ring (See Fig S10 for the definition of the plane) with the definition of the DNA bending angle *θ*. (E) The distributions of *θ* calculated from the simulations of the dimeric (SMC1/3 heads) and the tetrameric (SMC1/3 heads, STAG1, and NIPBL) complexes. (F) The initial structures of the simulations of the tetrameric complex with 20 bp DNA. Surface amino acid particles forming the DNA-gripping patch group are colored in blue. DNA model is colored in red and green.

As mentioned, in cryo-EM structure of the human cohesin-DNA complex, SMC1 and SMC3 have consecutive missing residues in their coiled-coil regions, and RAD21 has most of its IDR missing. The structures of these regions were predicted using AlphaFold Colab (30). Then, we sought to build a residue-resolution structural model of the whole human cohesin complex in the gripping state from the subunit structures of SMC1 (full length), SMC3 (full length), STAG1 (HEAT repeats domain, Residues 86-1052), NIPBL (HEAT repeats domain, Residues 1193-2628) and RAD21 (full length). To obtain the gripping state structure, we performed steered MD simulations to assemble 1. RAD21, and 2. [NIPBL and STAG1] to a SMC1/SMC3 dimer structure in consecutive simulations (Methods). Subunits are driven to their bound state by artificial harmonic bonds created based on selected inter-subunit contacts in the gripping state cryo-EM structure (14). As we approach the desired gripping state, the residue distances have become sufficiently small for the native contact potential constructed from the cryo-EM structure to take effect. At that stage, we removed artificial harmonic bonds (Fig S9). The DNA binding patches predicted by the simulations of each subunit were mapped on the resulting structural model of the whole human cohesin complex (Fig 5A).

Our structural model of the whole human cohesin complex showed the predicted DNA binding sites potentially involved in DNA loop extrusion (Fig 5A). Previous studies proposed various mechanisms to explain loop extrusion. One of them is the DNA segment capture model (13,31,32). This model describes the ATP hydrolysis cycle driving loop extrusion as three key steps: 1) ATP binding to the SMC1/3 heads induces head engagement and subsequent opening of the coiled-coil arms emanating from it. 2) A DNA segment (the increment of the main extruded loop in each cycle) is spontaneously inserted between the opened arms, with the two DNA ends held by the regions around the hinge and heads (Fig 5B). 3) After ATP hydrolysis and ADP release, the coiled-coil arms zip to close again while the DNA segment held around the hinge is pumped to the head area. Consistent with this model, we predicted strong DNA binding sites on the top surface of the SMC1/3 head dimer (C2, C3 in Table 1) and the hinge dimer (F1 in Table 1).

NIPBL binds to and quickly dissociates from the cohesin complex during loop extrusion and is proposed to stimulate ATP hydrolysis (11,33). Therefore, NIPBL is likely involved in step 2, after head engagement and before the ATP hydrolysis happens, with the spatial organization of SMC1, SMC3, NIPBL, STAG1, and the DNA like our whole complex based on the cryo-EM structure (14). In this stage, the distance between the hinge dimer and head dimer is around 10 nm (in the case where the coiled-coil arms of SMC1 and SMC3 are not folded, the distance is around 40 nm(34)), smaller than the persistence length of DNA of 45 nm (35). However, the loop extrusion speed (0.5–1 kbp/s) and ATPase rate (2 ATP/s) gives a rough estimation of step size up to ∼1 kbp (340 nm) (7,8). A more direct step size measurement of condensin, a member of the SMC complexes family with a similar size as cohesin, captures step sizes over 500 bp (170 nm) (36). The interaction between cohesin subunits and DNA held by the SMC1/3 head dimer in step 2 may help to rationalize this large step size through the bending of DNA at the top of SMC head dimer, which would allow DNA to deeply insert into the coiled-coil ring, resulting in a more extended DNA segment between hinge and heads (Fig 5B bottom).

Our whole complex structural model shows a group of DNA binding patches near SMC1/3 heads (C2, C3, H2, G1, G3 in Table 1). This group encircles the space where DNA is observed in the gripping state. The potential role of this patch group in step 2 of the DNA segment capture model is holding DNA on the top surface of SMC1/3 heads while bending DNA to favor the stabilization of a more extended DNA segment captured between the hinge and heads. The cryo-EM structure of the cohesin pentamer, including SMC1, SMC3, RAD21, and NIPBL, in the gripping state (14) identified DNA binding sites that largely overlapped with our predictions based on the simulations of the individual subunits. Interestingly, our simulations predicted one more site on STAG1 (G3). In the cryo-EM structure, the segment of DNA expected to bind to this site cannot be resolved, possibly due to structural flexibility around this region and the need to average over many complexes found in the same conformation in the cryo-EM structure.

We therefore sought to confirm the potential role of the complete patch group, including G3 on STAG1, in DNA loop extrusion. To this aim, we performed MD simulations of the cohesion complex consisting of SMC1/3 heads and emanating partial coiled-coil arms, STAG1, and NIPBL with a 78 bp DNA fragment [the same length and sequence as in the cryo-EM structure (14)] manually placed as in the cryo-EM structure (in which only 17 bp out of the 72 bp can be resolved, Fig 5C, top). In the simulation trajectories, we observed that DNA sharply bends as it reaches the equilibrium (Fig 5C top, Movie 2). We measured the angle of DNA bending projected on the plane normal to the plane spanned by SMC1/3 coiled-coil arms and aligned along the bisector defined by the coiled-coil axis (Fig 5D, Fig S10 and Methods for details) The mean angle of DNA bending was 32.4°±0.3°. We also performed the simulations without STAG1 and NIPBL (Fig 5C bottom) and found that the mean angle was reduced to 5.2°±0.8° (Fig 5E). The larger angle found in the system with the full DNA-gripping patch group is expected to facilitate the stabilization of a larger loop captured between the heads and hinge and therefore a larger step size of loop extrusion. When binding to the DNA-gripping patch group, DNA stayed in contact with at least part of the patch group in all 20 simulation trajectories (Fig 5C top). On the contrary, when DNA was placed on the SMC1/3 head dimer without STAG1 and NIPBL, dissociation from the top surface was observed in 16 out of 20 trajectories (Fig 5C bottom, Movies 2 and 3). More stable DNA binding in the presence of the patch group in the whole complex might facilitate extrusion by preventing the slippage of the captured DNA segment (which would lead to an unproductive ATP cycle).

To investigate the cooperative contribution of the predicted binding sites to affinity, we also performed MD simulations to quantify the binding strength of the group. A tetramer consisting of SMC1/3 heads and emanating partial coiled-coil arms, STAG1, and NIPBL was simulated with a 20 bp DNA molecule. Unlike previous systems A to H, which all have relatively simple, close-to-globular structures, the tetramer forms a tunnel structure, and the group of DNA binding patches is located on the inner sides of the tunnel. To increase the chance of DNA-protein interaction, we initially placed one DNA near the target patch group (Fig 5F). When treated as one patch, the dissociation rate constant of the group was 1.21 × 10^―7^, while the strength of its component patch C2 (on the top surface of the SMC1/3 head dimer), C3 (on the top surface of the SMC1/3 head dimer), H2 (on the neck of NIPBL connecting its N-terminal handle and U-shaped hook), G1 (on the inside surface of STAG1 U-shaped hook), G3 (on the inside surface of STAG1 U-shaped hook) were 1.49 × 10^―5^, 1.49 × 10^―5^, 9.32 × 10^―7^, 7.04 × 10^―6^, and 9.37 × 10^―6^, respectively. This marked decrease in the dissociation rate constant indicates the cooperative effect of multiple patches to stabilize DNA binding. Thus, these simulation results suggest that STAG1 and NIPBL facilitate loop extrusion by contributing to stable DNA binding and sharp DNA bending.

## Discussion

In this study, we employed residue-resolution MD simulations to identify strong DNA binding patches on the individual cohesin subunits, which may play an essential role in the molecular mechanisms of DNA loop extrusion. The validity of many of the identified patches was confirmed by comparison to mutagenesis experiments. We then constructed a model of the whole cohesin complex, including SMC1, SMC3, STAG1, NIPBL, and RAD21, which suggests how the DNA binding patches may cooperate during extrusion. Our findings provide vital clues for how the proposed DNA segment capture model of loop extrusion may be implemented in practice.

In particular, we used AlphaFold and steered MD simulations to model the ATP-bound state where SMC1/3 head domains are engaged, DNA is gripped at the top surface of the SMC1/3 head dimer, which most likely represents the state in which a DNA segment is captured by the ring compartment formed by SMC1/3 coiled-coil arms. In this state, we discovered the significant cooperative effect of multiple DNA binding patches in stabilizing and sharply kinking DNA near the top surface of the SMC1/3 head dimer, possibly facilitating the capture of a large loop (larger than the cohesin itself) to be extruded throughout the ATP hydrolysis cycle. Our study also revealed the importance of accessory protein subunits STAG1 and NIPBL. In their absence, DNA is unstable and likely to dissociate from cohesin, inhibiting productive translocation.

Although we included RAD21 in our model of the whole cohesin complex, we note that RAD21 has long intrinsically disordered regions, which were not observed by cryo-EM experiments. The path of RAD21 IDR in our model (Fig 5A green) is one possibility and should not be taken as definite. However, this model connects SMC1 and SMC3 head domains with a full-length RAD21 model, forming a topologically closed ring structure, and can be used as a starting point to study the behavior of the cohesin complex following the DNA segment capture step: when the SMC1/3 head disengages, RAD21 IDR should move with considerable flexibility. At the same time, its N-terminus and C-terminus stay bound to SMC3 and SMC1, respectively. RAD21 is expected to restrict the DNA loop extruded by cohesin, preventing loop dissolving when SMC1/3 heads disengage. While RAD21 was not considered in our analysis of the DNA binding patches due to its highly disordered nature, future studies should address how RAD21 may be able to modulate DNA binding to the whole cohesin complex.

According to the DNA segment capture model, after SMC1/3 head disengagement, coiled coils of SMC1/3 should “zip up” from hinge to head, pushing the DNA segment captured between hinge and head into the head area, resulting in the expansion of the DNA loop captured in the SMC1/3 head dimer-RAD21 compartment. This process can be studied using our model as a starting point, using longer DNA (larger than 1 kbp). However, to simulate the closing process of SMC1/3 coiled coils, experimental information on interactions between SMC1/3 coiled coils in their closed state (also known as APO state or I shape state) is required. Such data exist for yeast cohesin but not human cohesin considered in the current work.

Although we have discussed the potential roles of the identified DNA binding patches in the DNA segment capture model, our study does not rule out the possibility of other loop extrusion models. The swing-and-clamp model (9) proposes that DNA translocation is realized in two steps: 1) the SMC1/3 coiled coils extend, and the hinge dimer binds to downstream DNA. 2) Coiled coils fold, bringing the hinge dimer close to the head dimer and handing over downstream DNA from the hinge domain to the head region. Our study identified a strong DNA binding patch at the hinge dimer (F1 in Table 1), indicating that it can capture DNA. The swing-and-clamp model also suggests that NIPBL can dissociate from the head region during step 1, bind to the hinge domain, and facilitate searching for downstream DNA [the NIPBL-hinge proximity is indicated by Förster resonance energy transfer (FRET) experiments (9), while NIPBL searching and binding downstream DNA remains without direct experimental evidence]. Our study identified a strong patch at the “nose” region of NIPBL (H1 in Table 1). This patch has one order of magnitude stronger affinity than the patch on the hinge dimer and two orders of magnitude stronger affinity than the patches on the top surface of the head dimer. In the DNA gripping state, this strong patch is covered due to NIPBL-STAG1 binding, but when NIPBL dissociates from cohesin [a behavior confirmed by *in vitro* (7) and *in vivo* experiments (11)], this patch should be considered a candidate for the search of free DNA in step 2. Our study adds some credibility to the hypothesis made in the swing-and-clamp model and provides a possible mutation site for future experiments to verify this hypothesis.

Finally, several loop extrusion models assume the existence of a DNA anchor site to explain how DNA translocation may be converted into loop extrusion. Accordingly, one end of the DNA loop is translocated through mechanisms such as DNA segment capture or swing-and-clamp, while the other end remains anchored at a specific cohesin site. This anchor may be realized by DNA binding or encircling by RAD21 and preventing it from diffusing. The strong DNA binding patch we identified on the emanating coiled coils from the SMC1 head domain (A1) is also a candidate site for a DNA anchor.

## Materials and methods

### Modeling of the subunits and the whole cohesin complex

AlphaFold 2 (30) was employed to create the full length model of the cohesin subunits SMC1 and SMC3 (see below). These models were later used as a starting point for steered MD simulations to model the complete cohesin complex. AlphaFold 2 tends to favor compact, globular structures even when predicting the long antiparallel coiled-coil structures in SMC proteins (AlphaFold Protein Structure Database entry: Q14683, Q86VX4). To generate a model with realistic extended coiled coils, we predicted, using AlphaFold Colab (30), only the parts of coiled-coil structures missing from the cryo-EM structure (14). The amino acid sequence of SMC1: residues 203-490, 656-1030, and SMC3: residues 243-492, 684-926 was used as input. This yields reasonable structures as evaluated by pLDDT scores and compared with available crystal structures of their homologs (budding yeast SMC1 and SMC3, PDB ID: 7OGT, Fig S11). Non-helical regions at the middle part of coiled coils in 7OGT were defined as “elbows” where coiled coils fold. Elbow position in human SMC1 and SMC3 can be predicted by finding the homologous sequences of yeast elbow. Sequences of SMC1 and SMC3 of human, mouse, budding yeast, and fission yeast were aligned using the multiple sequence alignment server T-coffee (37). AlphaFold predicts the same positions as the folding point of coiled coils.

We used MODELLER (38) to construct a full-length molecule of SMC1 using head and hinge domain structures from the cryo-EM structure (PDB ID: 6WG3) (14) and AlphaFold modelled coiled-coil structures as templates. A short part of the coiled coil connected to the head domain is observed in 6WG3, which also exists in the coiled coils modelled by AlphaFold. This overlap prevented unnatural angles when MODELLER tried to connect two parts. In the hinge domain observed in 6WG3, the N terminus of the available structure starts with an alpha helix, continuous with an upstream alpha helix in the coiled-coil region. Using this as a restrain, MODELLER connects the coiled coils with the hinge domain without producing unnatural angles. The resulting full-length SMC1 model has the all-atom resolution.

Similar to SMC1, the cryo-EM structure of SMC3 also captures a short coiled-coil part connected to the head domain and an alpha helix in the hinge domain continuous with the coiled coils (14). Using restraints similar to those of SMC1, the full-length SMC3 was modelled using MODELLER.

We modelled the structured regions of STAG1 (residues 85-1052) and NIPBL (residues 1130-2628) using MODELLER, which generates flexible loop structures to complete the loops missing in 6WG3.

### DNA binding simulation assays

Each target system (SMC1 head domain; SMC3 head domain; SMC1/3 head dimer; SMC1 hinge domain; SMC3 hinge domain; SMC1/3 hinge dimer; STAG1; NIPBL; DNA gripping tetramer containing SMC1 head domain, SMC3 head domain, STAG1 and NIPBL) was separately simulated with short duplex DNA fragments. For SMC1 and SMC3, we simulated a stand-alone head or hinge domain with DNA instead of the full-length molecule. We also simulated the head dimer and the hinge dimer as their bound state in 6WG3 (14).

Each simulation contains B-type DNA fragments with the same sequence, ATAGTGATTATGAAAACTTT [a randomly generated 20 bp sequence with the same GC content as the DNA sequence used in cryo-EM (14)]. For small systems (SMC1 hinge domain; SMC3 hinge domain) one DNA fragment is used. For larger systems (SMC1 head domain; SMC3 head domain; SMC1/3 head dimer; STAG1; NIPBL), three DNA fragments were used to increase binding site search efficiency. For the DNA-gripping tetramer, one DNA fragment was manually placed near the DNA-gripping patch group to induce efficient DNA-protein interaction. The B-DNA structure of the fragment at all-atom resolution was generated using the DNA Sequence to Structure tool (39).

CafeMol generated the residue-resolution models of the various systems based on the AICG2+ (25) and 3SPN2.C (20) models for proteins and DNA, respectively. Each amino acid is represented by one bead centered at the C_α_ atom, while each nucleotide is represented by three beads corresponding to phosphate, sugar, and base groups. The AICG2+ model stabilizes a protein in its native state through bonded and non-bonded interactions based on the reference structure. Proteins and DNA interact via excluded volume and Debye-Hückel electrostatic interactions. Ionic strength was set at 300mM, higher than physiological conditions, to speed up the dissociation between DNA and a protein. In our simulations, the charges of surface beads in the residue-resolution protein models were optimized by RESPAC (26) to match the surface electrostatic field calculated from the respective all-atom structures using the PDB2PQR and APBS software tools (40). The combination of AICG2+ and 3SPN2.C models has been successfully applied to study the dynamics of many protein-DNA systems (21,22,41), as it allows a significant speedup compared to all-atom models while retaining sufficient detail to reveal critical insights into molecular mechanisms.

Simulations for SMC1 head domain, SMC3 head domain, head dimer, STAG1, and NIPBL were carried out in a 200 Å each side cubic box with periodic boundary. The SMC1 hinge domain, the SMC3 hinge domain and the hinge dimer were simulated in a 100 Å each side cubic box with periodic boundary. The DNA-gripping tetramer was simulated without setting a boundary. Each system was simulated using Langevin dynamics for 20 trajectories (each 10^8^ steps of *Δt* = 0.15*fs*). Note that CafeMol uses a low friction parameter to speed up the dynamics, and we used an ionic strength at 300mM, higher than the physiological condition to speed up the dissociation of DNA-protein binding, so the protein-DNA interaction time scale directly obtained from the simulations does not represent the actual time scale *in vivo*.

### DNA binding-patch calling and dissociation rate constant calculation

A residue was considered in contact with a DNA fragment when the minimal distance between the residue and any atom from the DNA fragment was below a threshold (set to 8Å, slightly larger than the Debye length of 5.6Å). Trajectories were analyzed using the Python library MDanalysis (42).

Surface residues were clustered into binding patches according to Jaccard distance that describes how often residues bind to the same short DNA fragment (43) (Fig S3):

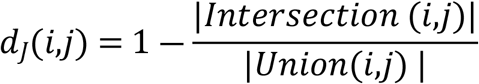

where the second term describes similarity between two sample sets, *Intersection* (*i,j*) is the number of trajectory frames in which residues *i* and *j* are in contact with the same DNA molecule, and *Union*(*i,j*) is the number of frames in which either residue *i* or *j* is in contact with any DNA fragment. Residues were hierarchically clustered (44) into binding patches using their Jaccard distance. Jaccard distance calculation and clustering were done using R (45).

After identifying binding patches, a binding event was considered a continuous series of timesteps where at least one residue in the binding patch of interest was in contact with a DNA fragment of interest. For each binding patch, from the distribution of the lifetime of all binding events for each DNA fragment in all trajectories, the survival probability of binding events was calculated as 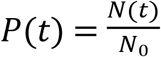, where *N*(*t*) = ∑ _τ>*t*_ *n* _*t*_ is a number of events with a lifetime larger than *t*, and *N*_0_ is the total number of events. The survival probabilities displayed two regimes, one corresponding to transient interactions giving short lifetimes typically shorter than some breakpoint *t*_1_, and one corresponding to stable binding giving lifetimes larger than *t*_1_ (Fig S5). To estimate the dissociation rate *k*_*off,stable*_, we fit the In 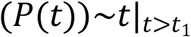 data points to a linear model. Discrete-time points {*t*_*i*_} were chosen so that {*N*(*t*_*i*_)} formed an arithmetic progression. In this way, data points can be given the same weight when performing linear regression. The breakpoint *t*_1_ was determined using the R package called “segmented” (46).

### Modeling of the whole cohesin complex in gripping state

Here, we describe the simulation protocol employed to generate the complete pentameric model of cohesin in the gripping state. We start from the full-length SMC1/3 complex generated by AlphaFold2 (Fig S9A and B), and we drive this toward the cryo-EM reference with PDB ID 6WG3 (14) by a combination of steered MD and switching-Gō simulation protocol (47). In the initial structure, SMC1 and SMC3 dimerize solely at the hinge but not at the heads, while the coiled coils folded at the two elbow positions. To allow the structure to relax toward the reference gripping state, the local bond length, bond angle, and dihedral angle interactions, and non-local contact interactions for the loop structures at the elbow positions of SMC1 and SMC3 were replaced by generic flexible local potentials (48). Intra-protein native contacts for positions except the elbow were based on the initial AlphaFold models. In contrast, SMC1/3 inter-protein native contacts were solely based on the hinge dimer of the reference 6WG3 structure in the first simulation phase.

We then set up a first steered MD simulation to relax the tightly folded initial structure so that the two subunits can interact as in the target 6WG3 gripping state (Fig S9A and B). The SMC1/3 hinge was softly fixed by anchoring one amino acid particle *A*_1_ to its initial position using a harmonic spring

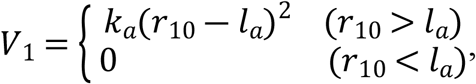

where *r*_10_ is the distance between the position of *A*_1_ and its initial position. Two ghost particles were defined at the position of two amino acid particles *A*_2_ and *A*_3_ in the SMC1 and SMC3 head domains, respectively. Ghost particles were connected to corresponding amino acid particles via a harmonic spring:

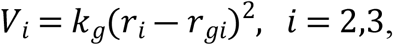

where *r*_*i*_ is the position of *A*_*i*_, and *r*_*gi*_ is the position of the ghost particles. Ghost particles move in a fixed direction at a constant velocity, pulling the head domains away from hinge domains (Fig S9A).

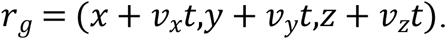

The steered MD simulation was performed for 10^6^ steps with *k*_*a*_ = 0.1*kcal* · *mol*^―1^ · Å^―2^, *k*_*g*_ = 1.0*kcal* · *mol*^―1^ · Å^―2^, and *v*_*g*_ = 0.001Å · (unit time)^―1^ (unit time in CafeMol is 49fs). This is then followed by an unbiased MD simulation of 10^7^ steps to obtain relaxed structures. The steered MD and the unbiased relaxation were conducted without setting a boundary. Simulation timestep and protein-protein interaction force-field were set as in the previous “DNA binding simulation assays” section.

Starting from the output of the previous simulations, we then proceeded to construct a model of the SMC1-SMC3-RAD21 trimer, in which SMC1 and SMC3 dimerized at the hinge, head domains were disengaged, and RAD21 connected the SMC1 head and the SMC3 head to form a closed ring (Fig S9C and D). We started from the extended SMC1/3 dimer structure and the structure of the relaxed full-length RAD21 generated by AlphaFold. AlphaFold predicted the majority of RAD21 as unstructured, with low pLDDT scores, indicating IDRs (30). Disordered regions predicted from sequence using DISOPRED3 (49) agreed with AlphaFold2 predictions. Based on the 6WG3 target, we manually picked five pairs of interacting amino acid particles, two from the SMC1-RAD21 binding interface and three from the SMC3-RAD21 interface. Harmonic bonds *V* = *k*_*b*_(*r*_*ij*_ ― *l*_0,*ij*_)^2^ were applied to each pair, with *r*_*ij*_ being the distance between two amino acids of each pair and bond length *l*_0,*ij*_ set as the distance of each pair in 6WG3. The steered MD simulations were performed first for 10^6^ steps with *k*_*b*_ = 0.001*kcal* · *mol*^―1^ · Å^―2^, and then for 10^6^ steps with *k*_*b*_ = 0.1*kcal* · *mol*^―1^ · Å^―2^. This successfully generated a SMC1-SMC3-RAD21 complex with a closed ring topology.

We then constructed a model of the SMC1-SMC3-NIPBL-STAG1-RAD21 pentamer (Fig S9E and F). To set up the initial structure for the steered MD simulation, we initially placed the STAG1 and NIPBL subunits near the SMC1-SMC3-RAD21 trimer structure obtained in the previous step. An analogous protocol as before was followed to add harmonic bonds between the STAG1-RAD21 and NIPBL-RAD21 interfaces based on the target cryo-EM structure (14). Steered MD simulations were performed for 10^6^ steps with *k* = 0.001*kcal* · *mol*^―1^ · Å^―2^, and then for 10^6^ steps with *k* = 0.1*kcal* · *mol*^―1^ · Å^―2^. RAD21 harmonic bonds from the previous step were also turned on, with *k* = 0.1*kcal* · *mol*^―1^ · Å^―2^.

Finally, we constructed the SMC1-SMC3-NIPBL-STAG1-RAD21 pentamer (the whole cohesin complex) in the DNA gripping state (Fig 9F to H), producing the complete model consistent with the observed cryo-EM structure (PDB ID: 6WG3) (14) but including all amino acids in the complex. The same protocol for adding harmonic bonds was applied to the SMC1 head domain-SMC3 head domain, SMC1 head domain-NIPBL, SMC3 head domain-NIPBL, and NIPBL-STAG1 interfaces. Steered MD simulations were performed first for 10^6^ steps with *k* = 0.001*kcal* · *mol*^―1^ · Å^―2^, and then for 10^6^ steps with *k* = 0.1*kcal* · *mol*^―1^ · Å^―2^. Harmonic bonds from the previous step were also turned on with *k* = 0.1*kcal* · *mol*^―1^ · Å^―2^. At this stage, the protein interfaces described above were close enough for the native AICG2+ contacts based on the reference cryo-EM structure to be effective. We, therefore, switched off the previous harmonic bonds and added the AICG2+ intra-molecule native contact interactions based on the reference gripping state structure (PDB ID: 6WG3) (14), except for those at the SMC1 hinge domain-STAG1 and SMC1 hinge domain-NIPBL interfaces. The potential-switching simulation was performed for 10^6^ steps. Next, we added harmonic bonds to SMC1 hinge domain-STAG1 and SMC1 hinge domain-NIPBL interfaces and performed steered MD simulations for 10^6^ steps with *k* = 0.001*kcal* · *mol*^―1^ · Å^―2^ and for 10^6^ steps with *k* = 0.1 *kcal* · *mol*^―1^ · Å^―2^. Harmonic bonds at other interfaces were not applied here, but AICG2+ interactions maintained the protein subunit bindings. Lastly, potential-switching to the native contact interactions at the SMC1 hinge domain-STAG1 and SMC1 hinge domain-NIPBL interfaces was performed for 10^6^ steps to obtain the final structure of the whole cohesin complex in the DNA gripping state.

All simulations in this section were performed using CafeMol, using the same settings described for the unbiased MD simulations.

### Modeling of cohesin tetramer containing SMC1/3 heads, STAG1 and NIPBL to study the DNA-gripping patch group

To construct the initial structures in Fig 5C and F, we used previously constructed models of SMC1/3 head dimer, STAG1, and NIPBL to build a tetramer containing the DNA-gripping patch group we are interested in. We added harmonic bonds to the SMC1 head domain-NIPBL, SMC3 head domain-NIPBL, and NIPBL-STAG1 interfaces following the same protocol as the previous section. The SMC1 head domain–SMC3 head domain interface was held together by the AICG2+ intra-molecule native contact interactions based on the gripping state structure.

To simulate the effect of DNA-gripping patch group and on long DNA, 3SPN.2C model of 72 bp dsDNA [sequence: TGGTTTTTATATGTTTTGTTATGTATTGTTTATTTTCCCTTTAA-

TTTTAGGATATGAAAACAAGAATTTATC, the same as in the cryo-EM experiment (14)] was manually inserted in the tunnel surrounded by patches of interest as an initial structure. To simulate the effect of SMC1/3 head-dimer as a control, we manually placed the same dsDNA near the top surface of an SMC1/3 head-dimer model. To measure the dissociation rate constant of the DNA-gripping patch group, one 20 bp dsDNA (same sequence as the previous DNA dissociation simulations) was placed at two different positions near the patch group to create two different initial structures.

### Definition of DNA bending angle θ

From models of the SMC1/3 head dimer or tetramer containing SMC1/3 heads, STAG1, and NIPBL (Fig 5C), we picked the particles B_1_(SMC1/S161) and B_2_(SMC3/T165) representing the base of SMC1 and SMC3 coiled coils, E_1_(SMC1/E202) and E_2_ (SMC3/N243) representing the end of emanating SMC1 and SMC3 coiled coils. The center of the top surface of the SMC1/3 head dimer C is defined as the geometric center of B_1_ and B_2_. The plane spanned by two coiled coils was defined by vectors 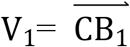 and 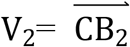. The plane normal to this plane was defined by the vector perpendicular to the plane *x* = V_1_ × V_2_ and the bisector *Z* = (V_1_ + V_2_)/2. The DNA trajectory is projected on the plane *xz* (Fig 5D).

From the coordinates of the DNA model projected on the plane *xz*, we selected the sugar particle of the first nucleotide (a_1_,b_1_) and the 11th nucleotide (a_2_,b_2_) at the 5’ end of each strand of the dsDNA. Considering the B-DNA helix is about 10 base pairs per turn, vectors 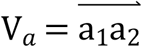 and 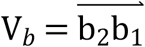 are approximately parallel to the track of the center of the DNA double helix. Therefore, we can calculate the DNA bending angle as 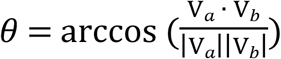, while not accounting for the effect of the natural twist of DNA helix. This twist adds a layer of complexity to the calculations, as it influences the overall shape and structure of the DNA.

## Acknowledgement

We would like to thank the members of the theoretical biophysics laboratory at Kyoto University for discussions and assistance throughout this work.

